# *In vivo* identification of apoptotic and extracellular vesicle-bound live cells using image-based deep learning

**DOI:** 10.1101/2020.04.14.040691

**Authors:** Jan Kranich, Nikolaos-Kosmas Chlis, Lisa Rausch, Ashretha Latha, Martina Schifferer, Tilman Kurz, Agnieszka Foltyn-Arfa Kia, Mikael Simons, Fabian J. Theis, Thomas Brocker

## Abstract

The *in vivo* detection of dead cells remains a major challenge due to technical hurdles. Here we present a novel method, where injection of fluorescent Milk fat globule-EGF factor 8 protein (MFG-E8) *in vivo* combined with imaging flow cytometry and deep learning allows the identification of dead cells based on their surface exposure of phosphatidylserine (PS) and other image parameters. A convolutional autoencoder (CAE) was trained on defined pictures and successfully used to identify apoptotic cells *in vivo*. However, unexpectedly, these analyses also revealed that the great majority of PS^+^ cells were not apoptotic, but rather live cells associated with PS^+^ extracellular vesicles (EVs). During acute viral infection apoptotic cells increased slightly, while up to 30% of lymphocytes were decorated with PS^+^ EVs of Dendritic cell exosomal origin. The combination of recombinant fluorescent MFG-E8 and the CAE-method will greatly facilitate analyses of cell death and EVs *in vivo*.

## INTRODUCTION

Billions of cells die every day in physiological and developmental processes (Nagata, 2018). Also, during viral infections cells are killed either directly by pathogens or by the immune system to limit pathogen expansion. However, despite high frequencies of cell death, it is extremely difficult to detect apoptotic cells *in vivo* (Surh and Sprent, 1994) due to lack of appropriate detection methods and highly efficient removal of dead cells by phagocytic macrophages (deCathelineau and Henson, 2003).

One hallmark of apoptotic cell death is the exposure of phosphatidylserine (PS) on the outer membrane surface of cells (Fadok et al., 1992; Martin et al., 1995). In addition to apoptotic cells, also extracellular vesicles (EVs) are PS^+^ (Hugel et al., 2005; Llorente et al., 2013; Martinez and Freyssinet, 2001; Thery et al., 2002; Wubbolts et al., 2003). EVs are very heterogeneous (Mathieu et al., 2019) and contain distinct nucleic acid, lipid and protein cargo derived from parental cells (Jeppesen et al., 2019). They may contribute to cell-to-cell communication and modulate physiological functions such as immunity, cancer progression, metastasis and transfer of viral genomes (Altan-Bonnet, 2016; Robbins and Morelli, 2014; Tkach and Thery, 2016). The concentration of EVs in bodily fluids can increase during cell death, cancer or infections (Altan-Bonnet, 2016; Robbins and Morelli, 2014). However, the major challenge to understand the role of EVs in biological processes is to study naturally occurring EVs *in vivo* as well as their target cells. This challenge remains unsolved, as specific reagents and analysis methods are lacking.

Fluorescently labeled Annexin V, which binds to PS, has been used to detect both, PS^+^ apoptotic cells and EVs (Heijnen et al., 1999). However, Annexin V requires elevated Ca^2+^-concentrations for PS-binding, which generates Ca^2+^-phosphate microprecipitates of EV-size, which can be mistaken for EVs (Larson et al., 2013). Furthermore, the Ca^2+^-requirement might make *in vivo* applications of Annexin V difficult and could interfere with many other downstream applications (van Engeland et al., 1998).

To reliably analyze PS^+^ EVs and dead cells *in vivo*, we have developed a recombinant PS-staining reagent by fusing Milk fat globule-EGF factor 8 protein (MFG-E8) (Hanayama et al., 2002) to enhanced green fluorescent protein (eGFP). MFG-E8 binds PS in a Ca^2+^-independent fashion with high sensitivity (Otzen et al., 2012) and already on early apoptotic cells (Shi et al., 2006). Furthermore, it binds to highly curved membranes (Shi et al., 2004), as those of small EVs.

Upon intravenous injection of MFG-E8-eGFP we performed imaging flow cytometry of fresh tissue cells on an ImageStream^x^ MarkII imaging cytometer, which allows detection of small particles with high sensitivity (Headland et al., 2014) and generates detailed images of individual cells (Pepperkok and Ellenberg, 2006). To automatically classify apoptotic vs. EV-decorated (EV^+^) cells, we developed a convolutional autoencoder (CAE) (Hinton and Salakhutdinov, 2006; Masci et al., 2011; Ranzato et al., 2007), which combines the advantages of traditional feature extraction (Blasi et al., 2016; Dao et al., 2016; Eliceiri et al., 2012) and deep learning (Eulenberg et al., 2017) for imaging flow cytometry. Using this pipeline, we show that MFG-E8-eGFP detects apoptotic as well as EV^+^ cells *in vivo*. In untreated mice EV^+^ hematopoietic cells are readily detectable at low frequencies *in vivo*. In contrast, irradiation or infection of mice with Lymphocytic choriomeningitis virus (LCMV) dramatically raised the frequencies of apoptotic and EV^+^ cells. Here, we analyzed B cells, DCs and T-cells among which we detected a striking increase of EV^+^ cells.

We provide a novel recombinant PS-binding molecule MFG-E8-eGFP, which, in combination with the deep learning CAE tool will give valuable information on the generation and function of EVs as well as on their target-cell specificities and will be most suitable to identify cell death *in vivo.*

## MATERIALS AND METHODS

### Mice

C56BL/6 mice were analyzed in sex and age-matched groups of 8–10 weeks of age. The SPF-status of the facility was tested according to the Federation for Laboratory Animal Science Associations (FELASA) recommendations. Animal experiment permissions were granted by the animal ethics committee of the Regierung von Oberbayern, Munich, Germany. All mice were bred and maintained at the animal facility of the Institute for Immunology, Ludwig-Maximillians-Universität München

### Generation of MFG-E8-eGFP

Murine MFG-E8-eGFP was produced from stably transfected HEK293 cells or purchased (#2002100; Bioconduct, France). Cells were grown in a Labfors Bioreactor (Infors, Switzerland) in serum-free medium (Ex-Cell 293, Sigma) for 5 days. Cells were removed from the cell culture supernatant (SN) by centrifugation (300g, 10min). 0.1% Triton-X 100 was added to solubilize membrane vesicles. SN was incubated under agitation for 1h. Debris was cleared by a high-speed centrifugation (40.000g, 60min) and filtration (0.2µm). MFG-E8-eGFP was then purified by FLAG affinity chromatography using M2-FLAG agarose beads (Sigma). Bound protein was eluted using an excess of FLAG peptide (Genscript, China). The eluate was concentrated using Sartorius spin columns with a cutoff of 30kDa (Sartorius). Lastly, MFG-E8-eGFP was further purified by gel filtration on an Äkta prime system with a Superdex 200 Increase 10/300 GL column (GE Healthcare).

### LCMV infections

LCMV Armstrong was propagated on L929 cells. Stocks were frozen at −80°C. For quantitation of virus titers focus-forming assays using Vero cells were performed as described previously (Pellegrini et al., 2011). For injections, viral stocks were diluted in sterile PBS. 2×10^5^ p.f.u. were injected intraperitonially per mouse.

### Preparation of single cells suspensions

Single cell suspensions of spleen and thymocytes were prepared by meshing organs through a 100µm nylon mesh. BM cells were flushed out from femur and tibia with PBS + 2%FCS using syringes. Erythrocytes were removed by centrifugation through a Pancoll cushion (Pancoll, PAN Biotech). Number of live cell was determined using a CASY cell counter (OMNI Life Science).

### FACS sorting of MFG-E8^+^ splenocytes and subsequent TEM

Single cell suspensions of splenocytes from LCMV infected, MFG-E8-eGFP injected mice were prepared by meshing organs through a nylon mesh and placed in PBS + 0.5% BSA. Erythrocytes were removed by centrifugation through a Pancoll cushion (Pancoll, PAN Biotech). Number of live cells was determined using a CASY cell counter (OMNI Life Science). Cells were stained with anti-CD45 APC, live/dead violet and anti-GFP FITC. After washing, cells were prefixed in 4% EM-grade PFA (Science Services) for 20 min before sorting. Cells were sorted on a FACSAriaIII (BDBiosciences) using a 130µm nozzle to keep shear forces to a minimum to avoid tearing off of the EVs. Cells were sorted into PBS + 0.5% BSA and pelleted at 300g. The cells were kept pelleted throughout all fixation, contrasting and infiltration steps. Cells were fixed for 15 min in 2.5% glutaraldehyde (EM-grade, Science Services) in 0.1 M sodium cacodylate buffer (pH 7.4) (Sigma Aldrich), washed three times in 0.1 M sodium cacodylate buffer before postfixation in reduced osmium (1% osmium tetroxide (Science Services), 0.8% potassium ferrocyanide (Sigma Aldrich) in 0.1 M sodium cacodylate buffer). After contrasting in 0.5% uranylacetate in water (Science Services), the pellet was dehydrated in an ascending ethanol series, infiltrated in epon (Serva) and cured for 48h at 60°C. Ultrathin sections (50nm) were deposited onto formvar-coated copper grids (Plano) and postcontrasted using 1% uranyl acetate in water and ultrostain (Leica). TEM images were acquired on a JEM 1400plus (JEOL) using the TEMCenter and tile scans with the ShotMeister software packages (JEOL), respectively.

### Imaging flow cytometry and data analysis

5×10^6^ cells were stained with appropriate antibodies for 20min on ice in PBS + 2% FCS and analyzed on an ImageStream^X^ MKII imaging flow cytometer (Merck). Mfge8-eGFP+ cells were gated using the IDEAS software. Then TIF-images of Mfge8-eGFP^+^ cells from each sample were exported (16-bit, raw) and analyzed by the CAE using a graphical user interface. The results were stored in two separated *.pop files containing the object numbers of apoptotic and EV-decorated cells. These object numbers were re-imported into IDEAS and two separate files containing only apoptotic or EV-decorated were generated. Next, from each sample, three files (one containing all cells, one containing only apoptotic cells and one containing only EV-decorated cells) were exported as fcs-files which were then further analyzed using FlowJo.

### Preparation of PKH26-stained EVs

40×10^6^ thymocytes were labelled with PKH26 red (Sigma Aldrich) according to the manufacturer’s protocol. Briefly, the cell suspension was washed with a serum-free DMEM medium (GIBCO) and resuspended in 1 ml of dilution buffer from the manufacturer’s labeling kit. The cell suspension was mixed with an equal volume of the labeling solution in the dilution buffer and incubated for 5 min at RT. Labeling reaction was stopped by addition of 2 ml fetal bovine serum (FBS) followed by with washing complete DMEM (10% FBS, 1% Penicillin). To induce apoptosis, cells were treated with 1μg/ml of Staurosporine (Sigma Aldrich) in serum free DMEM for 2 hours at 37°c followed by three washes. Cells were removed by centrifugation (500g). To collect PKH26-labeled vesicles, including apoptotic bodies, supernatant was ultra-centrifuged at 100,000g for 90 min. Prior to injection into mice, vesicles were resuspended in PBS.

### Data sets used for deep learning

All datasets examined in this study were acquired using the ImageStream^X^ MKII (Luminex). For the machine learning approach only brightfield images and MFG-E8-eGFP^+^ or PKH26^+^ fluorescent images were used. All images were cropped to 32×32 pixels and exported as 16-bit raw TIF images. No further pre-processing was performed on the pixel intensities (e.g. normalization or scaling). The *in vitro* annotated training dataset D1 consists of 27639 cells (27224 apoptotic, 415 EV^+^). The apoptotic cells in this dataset were stained with MFG-E8-eGFP *in vitro*, while the 415 EV^+^ cells were prepared from splenocytes after injection of PKH26-labeled vesicles. The *in vivo* annotated dataset D2 consists of 200 cells (100 apoptotic, 100 EV^+^). The M4 *in vivo* dataset consists of 382 cells (199 apoptotic, 183 EV^+^). The M1, M2, and M3 datasets were BM cells acquired from 3 irradiated mice and consist of 14922, 16545 and 17111 unannotated cells, respectively. The M5 and M6 datasets were acquired from BM of two non-irradiated mice and consist of 5805 and 5046 unannotated cells, respectively. Datasets D1 and D2 were imaged with a 40x objective, while datasets M1, M2, M3, M4, M5 and M6 were imaged with a 60x objective.

### Data analysis strategy

A novel pipeline combining unsupervised deep learning with supervised classification is used for cell classification, and compared to deep learning and classical feature based classification.

### Convolutional Autoencoder (CAE)

The CAE used in this study consists of a typical encoder-decoder scheme but with a channel-wise adaption: the encoder part is different for each input channel, while the decoder part of the network is used only during training, not for testing. The CAE was trained on 90% of M1 for 300 epochs, while the instance of the network that performed the best on the 10% validation set of M1 was saved and used for feature extraction in all subsequent experiments. The CAE consists of approximately 200,000 parameters and the exact architecture is shown in supplementary Figure S2. Each convolutional layer is followed by a batch normalization layer [batchnorm] and a ReLU activation [relu-glorot], with the exception of the last convolutional layer which is followed by a linear (activation) function (and no batch normalization). The mean squared error (MSE) of the reconstructed image was used as a loss function for training, whilethe mean absolute error (MAE) produced similar results in terms of classification accuracy. Adam [adam] was used to train the network, using a batch size of 64.

### Convolutional Neural Network (CNN)

The CNN used in this study for comparison is the exact same architecture as in (Eulenberg et al., 2017) and consists of approximately 3 million parameters. For comparison to the CAE, we also implemented a smaller version of the CNN architecture where each layer of the original architecture had 1/4 of the parameters, which resulted in a model with approximately 200 thousand parameters (same as the CAE). There was no significant difference between the performance of the original and downsized variants of the CNN in any of the experiments. As such, only the results of the original variant of the CNN are reported. This specific CNN architecture receives 64×64 images as input, while the available images are 32×32. As a result, all input images were padded with their edge values to fit the input dimension of the network. In all experiments the CNN was trained using Adam (Kingma and Ba, 2014).

### Cell-Profiler features

To compare to classical machine learning, the Cell-Profiler (CP) (Dao et al., 2016) pipeline from Blasi et al. (Blasi et al., 2016) was used for feature extraction. However, in our case the second channel corresponds to fluorescence intensity instead of darkfield.

### Random Forest

The scikit-learn (Pedregosa et al., 2011) Python implementation of the Random Forest (Breiman, 2001) algorithm was used. The number of trees (n_estimators) was set to 1000, while the number of features to assess at each split (max_features) was set to ‘sqrt’. In all subsequent experiments when we refer to CAE or CP accuracy, we mean the accuracy obtained by a Random Forest trained on the pretrained CAE (CAE-RF model) features or CP features (CP-RF model), respectively.

### CAE-RF/CP-RF

Both terms refer to a random forest trained on top of the features extracted using the pre-trained CAE introduced above or using Cell-Profiler, respectively. As such, training CAE-RF/CP-RF refers to training only the classification part of the method (RF).

### Confidence Intervals

Wilson’s method (Wilson, 1927) was used to calculate the proportion confidence intervals for classification accuracy.

### Code availability

The source code of this study is freely available at “https://github.com/theislab/dali”

## RESULTS

### MFG-E8-eGFP stains dying and PS^+^ live cells *in vivo*

To develop a robust, buffer-insensitive, fluorescent *in vivo* detection reagent for apoptotic cells, we fused murine MFG-E8 to enhanced green fluorescent protein (eGFP) for recombinant expression (Suppl. Fig. 1). The recombinant protein consisted of the full length MFG-E8 protein containing both C-domains (C1 and C2), of which especially the C2 domain confers PS-binding (Andersen et al., 2000). In addition, it contained the RGD-motif, which mediates binding to α_v_β_3_ integrin and facilitates phagocytosis of dead cells by macrophages (Hanayama et al., 2002). The purified MFG-E8-eGFP could identify similar frequencies of dying cells, as compared to the commercially available Annexin V *in vitro* (Fig. 1A). Double staining with both reagents showed that the same apoptotic cells bind Annexin V and MFG-E8-eGFP, when tested in a Ca^2+^-rich Annexin V-binding buffer (Fig. 1A). However, when conventional buffer was used, only MFG-E8-eGFP, but not Annexin V could detect apoptotic cells (Fig. 1A). This data indicates that MFG-E8-eGFP detects the entirety of dying cells similar to the reference reagent independently of specific buffer conditions.

**Figure 1.**
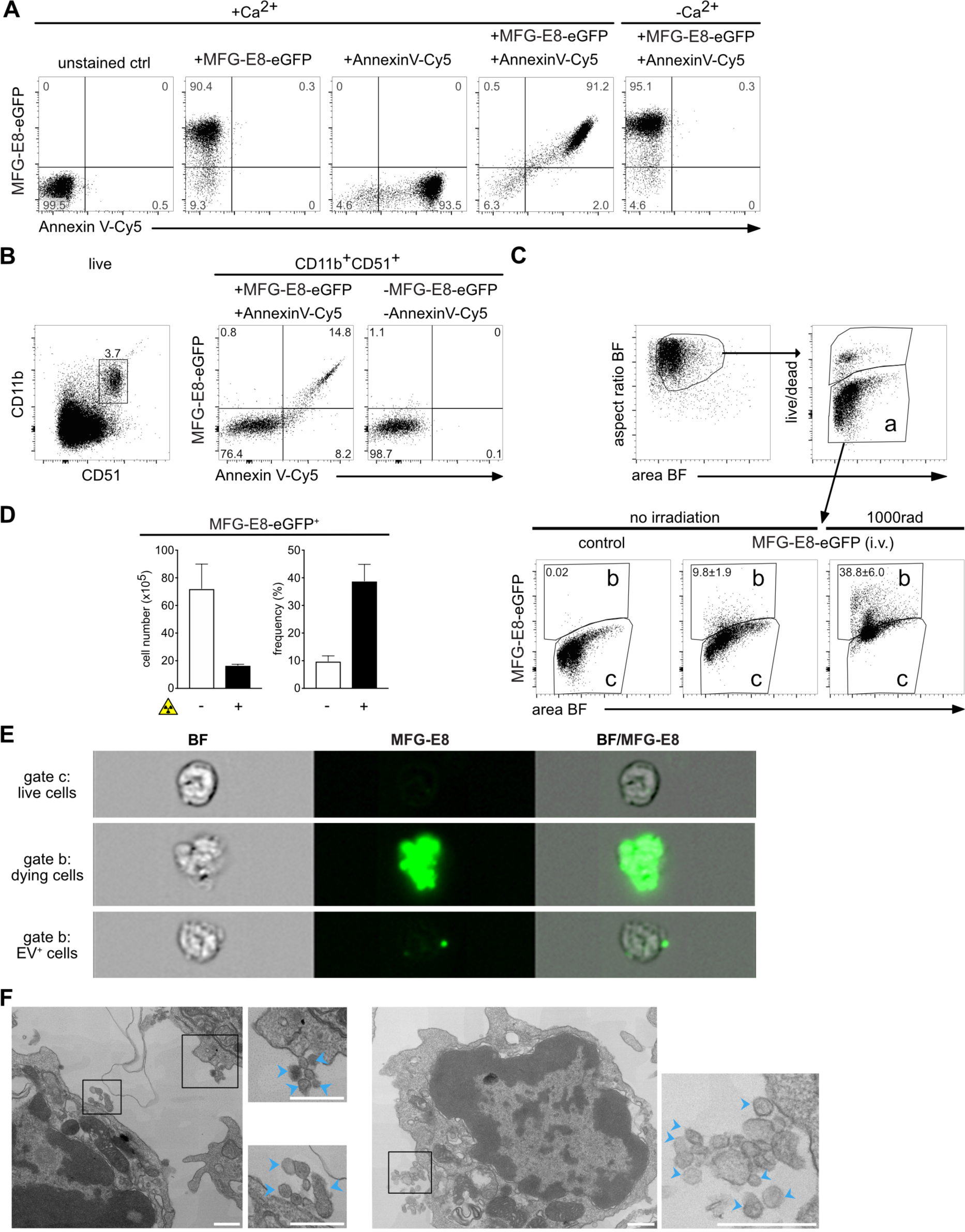
MFG-E8-eGFP stains apoptotic and EV^+^ cells *in vitro* and *in vivo.* (A) Staurosporine-treated (1µg/ml, 2h) apoptotic Jurkat cells were stained either with Annexin V-Cy5 or MFG-E8-eGFP or both reagents together in Ca^2+^-containing or Ca^2+^-free buffer as indicated. (B) To determine the degree of MFG-E8-eGFP binding by integrins via its RGD-motif by CD51 expressing cells, freshly isolated splenocytes were stained with MFG-E8-eGFP, AnnexinV, CD11b and CD51. Left dot plot shows gating of CD11b^+^CD51^+^ cells, middle plot shows MFG-E8-eGFP and Annexin V staining of CD11b^+^CD51^+^ cells, right plot shows unstained control. (C-D) To test MFG-E8-eGFP *in vivo* non-irradiated (n=3) and irradiated mice (1000rad, n=3) were injected with 100µg MFG-E8-eGFP i.v. 24h after the irradiation. 30min after the MFG-E8 injection mice were sacrificed and bone marrow (BM) cells were stained with live/dead violet and MFG-E8-eGFP followed by imaging flow cytometry on an Imagestream^X^ Mark II. Bar graphs display total numbers (left) and frequencies (right) of MFGE-E8^+^ cells with and without irradiation. Averages ± SD are shown. (E) BF and MFG-E8 images of live, MFG-E8^-^ (top), MFG-E8^+^ apoptotic (middle) and MFG-E8^+^EV^+^ (bottom) are shown. Scale bar: 7µm. (F) MFG-E8^+^ splenocytes were FACS-sorted (sorting strategy see Supplemental Fig. 2) and imaged by TEM. Two representative images of cells with attached extracellular vesicles (indicated by blue arrows) are shown. Scale bars: 500nm.

The RGD-motif present in MFG-E8-eGFP can potentially also bind to α_v_β_3_ and α_v_β_5_ integrins (Hanayama et al., 2002). To test if such binding might cause false-positive labeling of cells, we next stained spleen cells with anti-α_v_ (CD51) antibody, which mainly stained CD11b^+^ macrophages and monocytes (Fig. 1B, left panel). However, MFG-E8-eGFP only revealed CD11b^+^CD51^+^ cells, which were also Annexin V^+^ (Fig. 1B, right panel), indicating PS-specificity of MFG-E8-eGFP rather than binding via integrins.

Next, we administered MFG-E8-eGFP iv for *in vivo* labeling in order to avoid detection of artifacts generated during organ preparation, cell straining and other stress by *in vitro* handling. 30min after injection of MFG-E8-eGFP, bone marrow (BM) cells were harvested, stained *in vitro* with a viability dye (live/dead) to exclude necrotic cells (Fig. 1C, gate a). Due to their ruptured cell membranes we considered necrotic cells to be too damaged to extract reliable information and we focused only on life/dead^-^ cells in the live gate (Fig. 1C, gate a). Dying cells are rare in intact tissues due to their rapid removal (Henson and Hume, 2006). Accordingly, only approximately 10% of all BM cells in non-irradiated mice were MFG-E8-eGFP^+^ (Fig. 1C, gate b). To increase the rate of cell death, mice were γ-irradiated, which causes DNA damage and p53-mediated mechanisms of apoptosis within hours (Eriksson and Stigbrand, 2010). Therefore, while after irradiation the absolute numbers of live cells in the BM decreased due to cell death (Fig. 1D), the frequencies of MFG-E8-eGFP^+^ cells strongly increased (Fig. 1C,D). This indicated the general feasibility and specificity of an MFG-E8-eGFP *in vivo* application.

The individual images taken from cells within the live populations of Fig. 1C (gates a,b) showed that MFG-E8-eGFP^-^ cells had an intact rounded morphology typical for live cells (Fig. 1E). In contrast, MFG-E8-eGFP^+^ cells had cell bodies that were stained almost completely with MFG-E8-eGFP and showed densely stained apoptotic blebs indicating that cells are undergoing apoptosis (Fig. 1E, middle panel). However, within the same gate (Fig. 1E, gate b) we also found high numbers of cells that only showed very few, or even only one intensely stained MFG-E8-eGFP^+^ structure of subcellular size, whereas their cell body was unstained and had the rounded morphology of live, intact cells (Fig. 1E, lower panel). These particles were reminiscent of EVs and we next sorted MFG-E8^+^ lymphocytes from spleens for analysis by transmission electron microscopy (TEM). Attached to the sorted cells we could readily identify extracellular particles of 50-100nm diameter, a size typical for EVs (Fig. 1F). Therefore, MFG-E8-eGFP allows the analysis of apoptotic and PS^+^ EV-decorated live cells and we next set out to characterize them in more detail.

### An interpretable deep learning approach is able to discriminate EV-decorated cells from dying cells

The imaging analysis software IDEAS is very powerful in extracting and identifying features that help to discriminate different cell subsets (George et al., 2004). To generate a mask for separation of MFG-E8-eGFP^+^ cells into PS^+^ EV^+^ live or PS^+^ apoptotic cells, we manually selected 50 images of each type of MFG-E8-eGFP^+^ cells and analyzed their brightfield and fluorescence characteristics (Suppl. Fig. 3A). Based on the manually selected MFG-E8-eGFP^+^ apoptotic and EV^+^ cells we generated gates that included the majority of each cell type (Suppl. Fig. 3B). However, when we applied these definitions to cells without manual preselection in an unbiased fashion, these gates were insufficient to classify all events, leaving many cells uncategorized (Suppl. Fig. 3C).

To identify more reliable features for apoptotic cell discrimination from EV^+^ cells, we defined a “ground truth” as a basis for training different classification methods for cell sorting. For this, we generated EVs *in vitro*, fluorescently labeled them with PKH26 and injected these EVs into mice (Fig. 2A). Dead cells were defined using staurosporine treated thymocytes stained with MFG-E8-eGFP in addition to the apoptotic marker active caspase-8 (aCas8) (Ashkenazi and Salvesen, 2014) *in vitro* (Fig. 2B). We next tested three different machine learning approaches with these data: (i) a Convolutional Neural Network (CNN) for imaging flow cytometry (Eulenberg et al., 2017), (ii) our proposed method CAE-RF (a classifier trained on features learned by a CAE as displayed in the scheme of Fig. 2 (C,D) (iii) CP-RF, a classifier trained on pre-defined Cell Profiler features (Blasi et al., 2016). In order to estimate the effect of inter-experiment batch effects on classification performance, all methods were tested twice on the same *in vivo* stained dataset (manually selected apoptotic and EV^+^ cells from irradiated mice) and their performance was assessed using the Area Under of the receiver operating characteristic curve (AUC) (Hastie et al., 2009). During the first trial both, the training (MFG-E8-eGFP^+^ BM cells from irradiated mice) and test experiments were stained *in vivo*. In this case, all methods achieved near perfect classification performance of AUC>0.99 (Fig. 2E). During the second trial, the apoptotic cells of the training experiment were stained *in vitro* (Fig. 2B), introducing a batch effect for the classifiers to overcome (Fig. 2F). In this case, CAE-RF generalized the best (Fig. 2E; AUC=0.854), followed by the CNN (Fig. 2E; AUC=0.811), while CP-RF failed to generalize to the new experiment and was comparable to random guessing (Fig. 2E; AUC=0.508). CAE-RF was as accurate as a CNN in identifying apoptotic cells in a new experiment (Fig. 2E).

**Figure 2.**
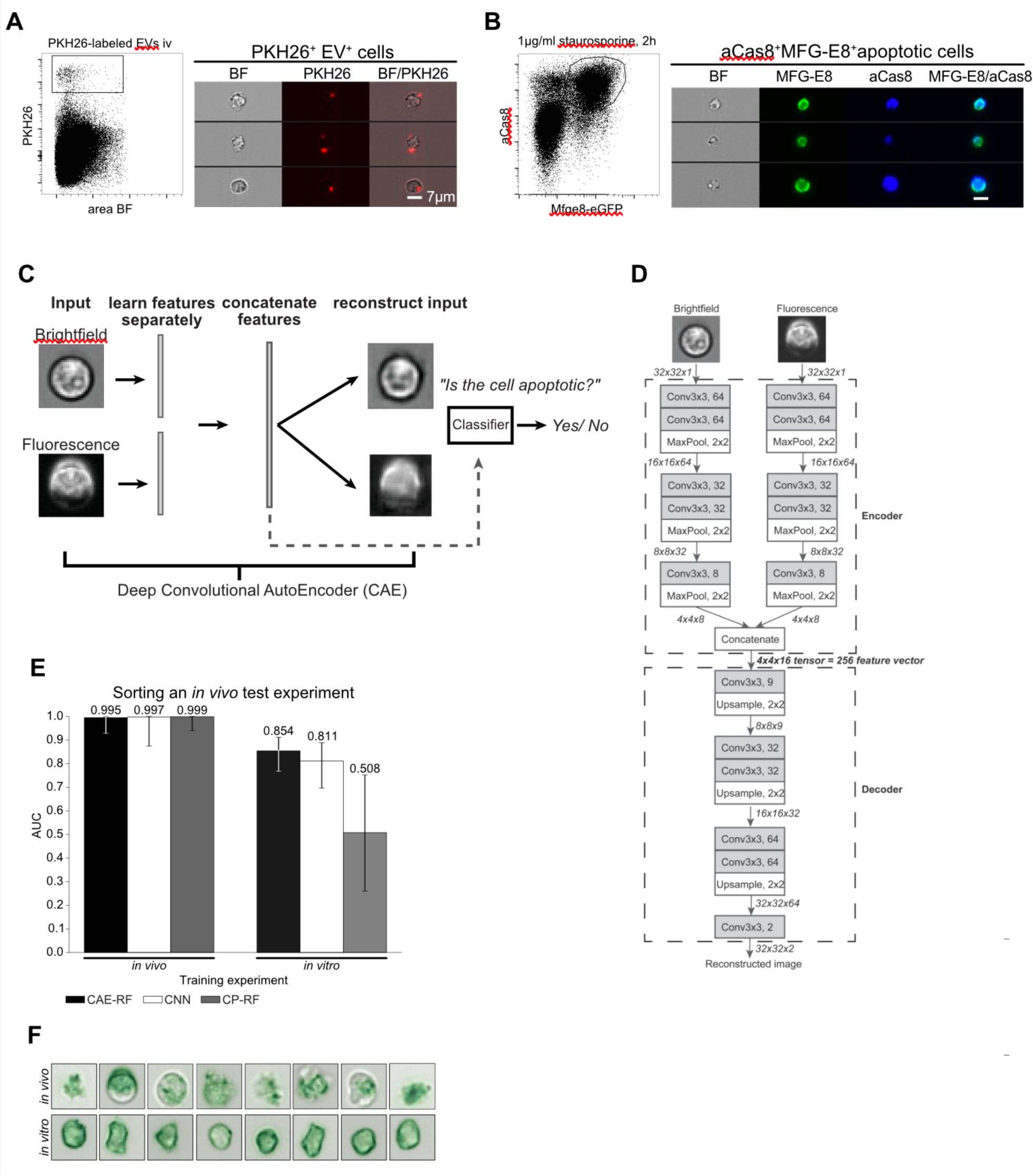
Discrimination of apoptotic vs. EV^+^ cell using machine learning. (A) To define a truth population for cells carrying EVs, PKH26-labeled EVs from staurosporine-treated thymocytes were injected i.v. into C56BL/6 mice. After 1h spleens were removed and analyzed by imaging flow cytometry. Images show splenocytes that are decorated with *in vitro* generated, PKH26-labeled EVs. (B) To define a truth population for apoptotic cells, staurosporine-treated thymocytes (1µg/ml for 2h) were stained with MFG-E8-eGFP (200ng/ml) and anti-aCas8 and analyzed by imaging flow cytometry. Images show MFG-E8-eGFP^+^aCas8^+^ apoptotic cells. (C) Using a Convolutional Autoencoder (CAE), both channels of the same image are separately encoded and then concatenated to form a 256-dimensional feature vector. During training features are learned in an unsupervised manner, by reconstructing the input images. In order to perform cell sorting, a classifier (Random Forest) is trained on a small subset of annotated cells. (D) Each input image consists of 32×32 pixels and 2 channels (Brightfield, Fluorescence). Every arrow in the figure corresponds to a data tensor. Each channel is encoded separately by an encoder network consisting of alternating convolutional and pooling layers. The encoder compresses each 32 ×32×1 (32×32 pixels, 1 channel) image into a 4×4×8 tensor. The encoded tensors of both channels are concatenated in a 4×4×16 tensor: the bottleneck of the CAE, which has a dual purpose. At training time, it is fed into the decoder part of the network which aims to reconstruct the input image in an unsupervised manner. At test time, the 4×4×16 bottleneck tensor is reshaped into the 256-dimensional feature vector of the input image that can be used in downstream tasks, such as classification of cell-subtypes. Each Convolutional layer performs 3×3 convolutions and is followed by a Batch Normalization layer and a ReLU activation function. The only exception to this rule is the last Convolutional layer (Conv3×3, 2), which is directly followed by Linear (identity) activation function. (E) Left: All models were trained on an *in vivo* stained dataset of 401 cells (M4), then tested on an independent *in vivo* stained dataset of 200 cells (D2). Right: the same models were trained on a new dataset of 27,639 cells (D1), where the apoptotic cells were stained *in vitro*, introducing a batch effect. Next, they were tested on the same 200 cell dataset as before (D2). Sorting performance is displayed as area under curve (AUC). (F) Demonstration of the batch effect introduced by *in vitro* staining of apoptotic cells. Right column: random subset of *in vivo* stained apoptotic cells. Left column: random subset of *in vitro* stained apoptotic cells. *In vitro* stained cells fluorescent staining is evenly distributed inside the cell, while *in vivo* stained cells exhibit more complex shapes and abnormalities in the distribution of the fluorescent dye. Error-bars correspond to 95% Wilson confidence intervals (n=200).

### For classification MFG-E8 fluorescence is more important than brightfield

The CAE-RF and CP-RF methods were trained using 5-fold cross validation (Hastie et al., 2009). Both CAE-RF and CP-RF agreed that features derived from the fluorescence channel were more important than brightfield features, for the task of apoptotic cell identification (Fig. 3A). Quantitative assessment of CAE-RF performance was done first by training on *in vivo* stained BM cells from irradiated mice. Then it was used to identify apoptotic and non-apoptotic cells in new experiments with data from irradiated (Fig. 3B; M2, M3) and non-irradiated mice (Fig. 3B; M5, M6). The performance of CAE-RF sorting was compared to standard gating on manually defined features (Fig. 3B). Both methods agree that more apoptotic cells are present in irradiated than in non-irradiated mice (Fig 3B). Subsequently, a subset of cells was manually annotated for each dataset, to quantitatively assess the classification performance of both methods CAE-RF (Fig. 3C) and manual gating using IDEAS features (Fig. 3D). In all cases, sorting with CAE-RF was more accurate than performing gating (Fig. 3C,D). Moreover, CAE-RF always characterized all cells, while manual gating resulted in some cells characterized as “unknown” since they did not correspond to any of the gates (Fig. 3D). Nonetheless, even if we discard the unknown cells from the calculation of classification accuracy (providing an advantage to gating), using CAE-RF was still more accurate (Fig. 3D).

**Figure 3:**
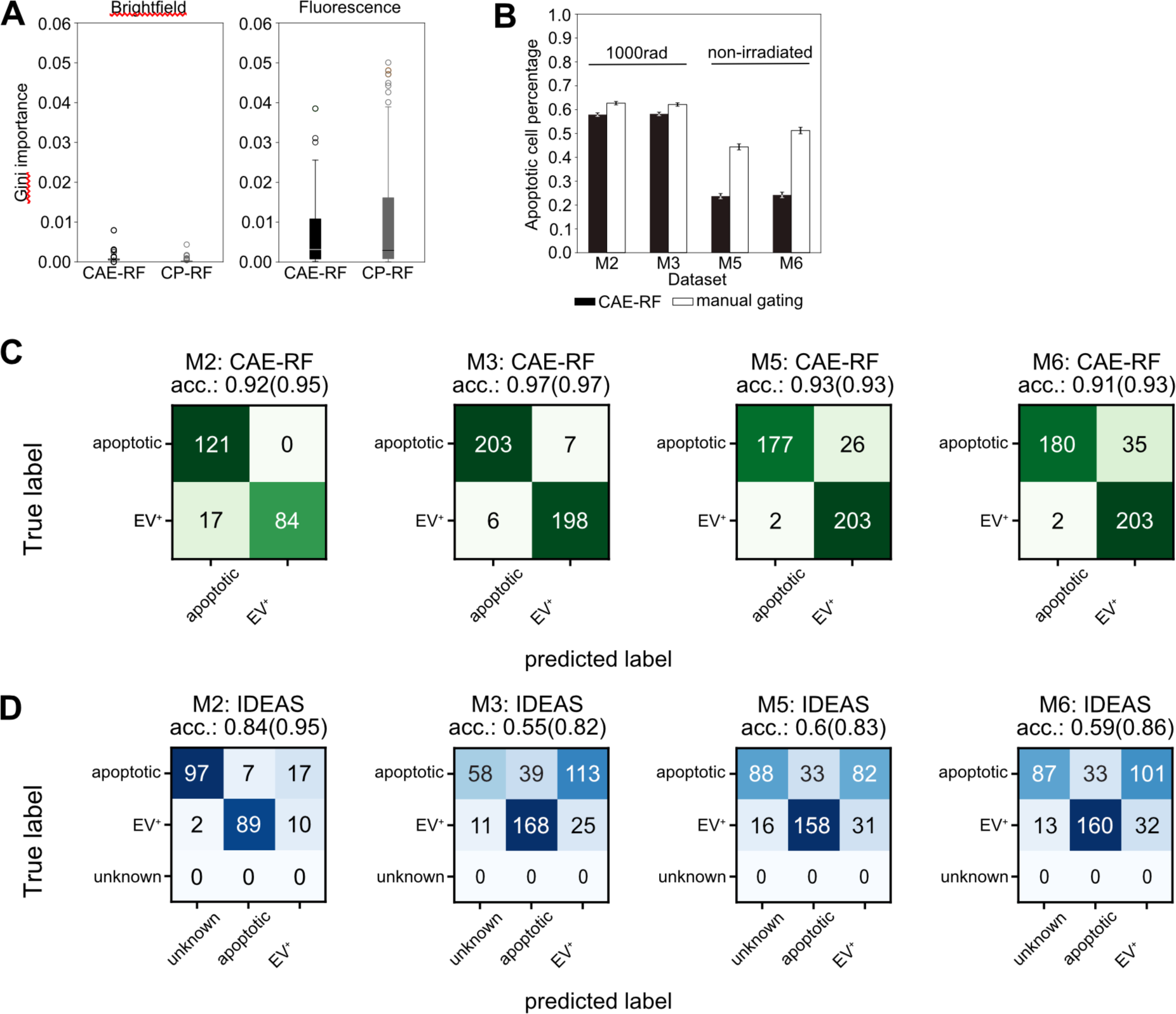
Convolutional Autoencoder performance. (A) Both CAE-RF and CP-RF identify the fluorescence channel (FL) features as more important than brightfield features (BF) for the task of apoptotic cell detection. Each boxplot visualizes the Gini importance of features belonging to the corresponding channel (FL or BF), as calculated by the random forest for each feature extraction method (CAE or CP). Fluorescence features have larger values of Gini importance than BF features. (B) When predicting on new non-annotated data, both CAE-RF classification and manual gating on IDEAS features predict more apoptotic cells in irradiated mice (M2, M3) and more cells with attached vesicles in healthy mice (M5, M6). Error bars correspond to 95% Wilson confidence intervals (n_M2_=16545, n_M3_=17111, n_M5_=5805, n_M6_=5046). A subset of cells was annotated manually for each dataset and sorting was performed using the CAE-RF (C) and IDEAS gating (D). “Unknown” cells fail to lie on the apoptotic or EV^+^ gate using IDEAS gating. The classification accuracy reported in parentheses for each confusion matrix corresponds to the accuracy if “unknown” cells are omitted.

### True dead and live EV^+^ cells can be sorted automatically by the novel deep learning approach

To challenge the accuracy of the CAE classifier, we next submitted image information from BM cells after irradiation to CAE-RF based sorting. As expected, the frequencies of live MFG-E8-eGFP^+^ cells strongly increased upon irradiation (Fig. 4A, B). Here, only cells were included that were live/dead^-^ apoptotic cells with intact cell membrane. MFG-E8-eGFP^+^ dying apoptotic cells had a higher MFG-E8-eGFP median fluorescence intensity than EV^+^ cells and both types could be clearly separated from each other by using CAE-RF (Fig. 4A). Although the rates of both, apoptotic as well as live EV^+^ cells increased upon irradiation, the EV^+^ cells were by far more frequent than dying cells (Fig. 4B). We analyzed both cell types further and used antibodies to CD11b^+^ myeloid cells and B220^+^ B lymphocytes, as prominent representatives of BM populations (Fig. 4C). As expected, we could detect only few B220^+^ B cells in irradiated mice (Fig. 4C), but a high frequency of them was classified as apoptotic (Fig. 4D, upper panel). This reflected the high sensitivity of B cells to γ-irradiation (Heylmann et al., 2014). In contrast, total numbers of myeloid CD11b^+^ cells were less decreased, as they have lower sensitivity to γ-irradiation (Rodrigues-Moreira et al., 2017) (Fig. 4D). However, also in the CD11b^+^ myeloid compartment, the frequencies of dying cells increased significantly upon irradiation (Fig. 4C, D). Frequencies of EV^+^ live cells also increased significantly in both, CD11b^+^ and B220^+^ cells during irradiation (Fig. 4C, D). Taken together, the CAE-RF module is able to reliably separate EV^+^ dying cells from PS^+^ live cells and can assist in more precise analyses of MFG-E8-eGFP^+^ cells.

**Figure 4.**
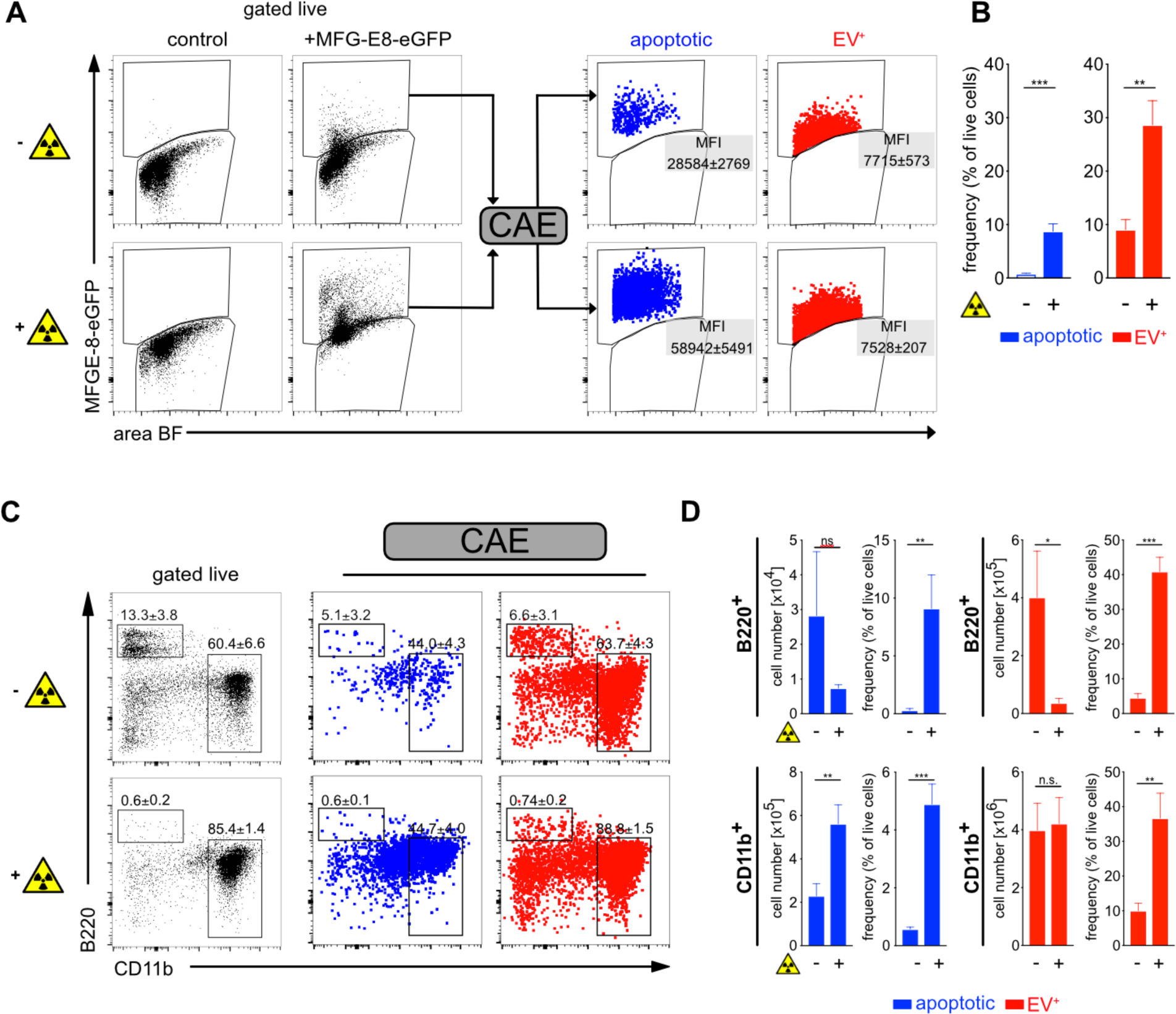
Using deep learning to discriminate apoptotic and EV^+^ cells. Non-irradiated controls (n=3) and lethally irradiated mice (1000rad, n=3) were injected with 100µg MFG-E8-eGFP i.v. 24h after the irradiation. 30min later mice were sacrificed. Bone marrow (BM) cells were analyzed by imaging flow cytometry. (A) To identify apoptotic and EV^+^ cells, cells were analyzed using IDEAS, CAE-RF and FlowJo. First, single cells were gated using the brightfield (BF) aspect ratio and the area of the BF signal. Then necrotic cells (live/dead^+^) were excluded from further analysis (Suppl. Fig. 5A). MFG-E8-eGFP^+^ cells were gated and their TIF images (16-bit, raw) exported using the IDEAS software. CAE-RF results with the classification apoptotic/EV^+^ were re-imported into IDEAS and separate fcs-files containing all cells or only MFG-E8-eGFP^+^/apoptotic cells and MFG-E8-eGFP^+^/EV^+^ cells were generated for further analysis in FlowJo. Apoptotic (blue) and EV^+^ (red) cells are shown in dot plots and their MFI of the MFG-E8 signal is displayed. (B) Bar graphs show apoptotic (blue) and EV^+^ (red) cells as frequency of live cells in non-irradiated and irradiated mice. Averages ± SD are shown. (C) Left dot plots show B220 and CD11b stained BM cells, gated on non-necrotic live/dead^-^ cells. Middle and right dot plots show B220 and CD11b expression of MFG-E8-eGFP^+^ cells classified by the CAE as apoptotic (blue) or EV^+^ (red), respectively. (D) Bar graphs show total numbers and frequencies of B220^+^ and CD11b^+^ apoptotic (blue) and EV^+^ (red) cells in non-irradiated and irradiated mice. Bar graphs show means ± SD, n=3.

### Distinction of dying from EV-decorated cells during acute infection with LCMV

Having developed a reliable automated method to discriminate apoptotic from EV^+^ cells, we next analyzed spleens of mice during an LCMV infection, which is known to induce cell death during the acute infection phase (Matter et al., 2006). Here, cell death is mainly caused by innate and adaptive immune mechanisms (Borrow et al., 1995; Matter et al., 2006; Odermatt et al., 1991). LCMV infection caused a strong increase in frequencies and total numbers of live MFG-E8-eGFP^+^ cells (Fig. 5A). To differentiate apoptotic from EV^+^ live cells we CAE-sorted their images. This revealed that both, apoptotic as well as EV^+^ cells significantly increased upon LCMV infection (Fig. 5B). However, the frequencies of PS^+^ live cells were more than 10-fold higher as compared to those of apoptotic cells, before as well as during an LCMV infection (Fig. 5B). More detailed analysis showed that the highest numbers of dying cells in non-infected mice were present within the CD19^+^ B cell and CD19^-^TCRβ^-^ non-B/T cell populations (Fig. 5C). Upon LCMV infection, both, dying and live EV^+^CD19^+^ B cell and CD19^-^TCRβ^-^ non-B/T cells further increased, but live EV^+^ B cells outnumbered dying B cells approximately 10-fold (Fig. 5C). In addition, especially CD8^+^ T cells showed increased frequencies of apoptosis and EV-decoration upon LCMV infection (Fig. 5C). We next analyzed individual populations in more detail.

**Figure 5.**
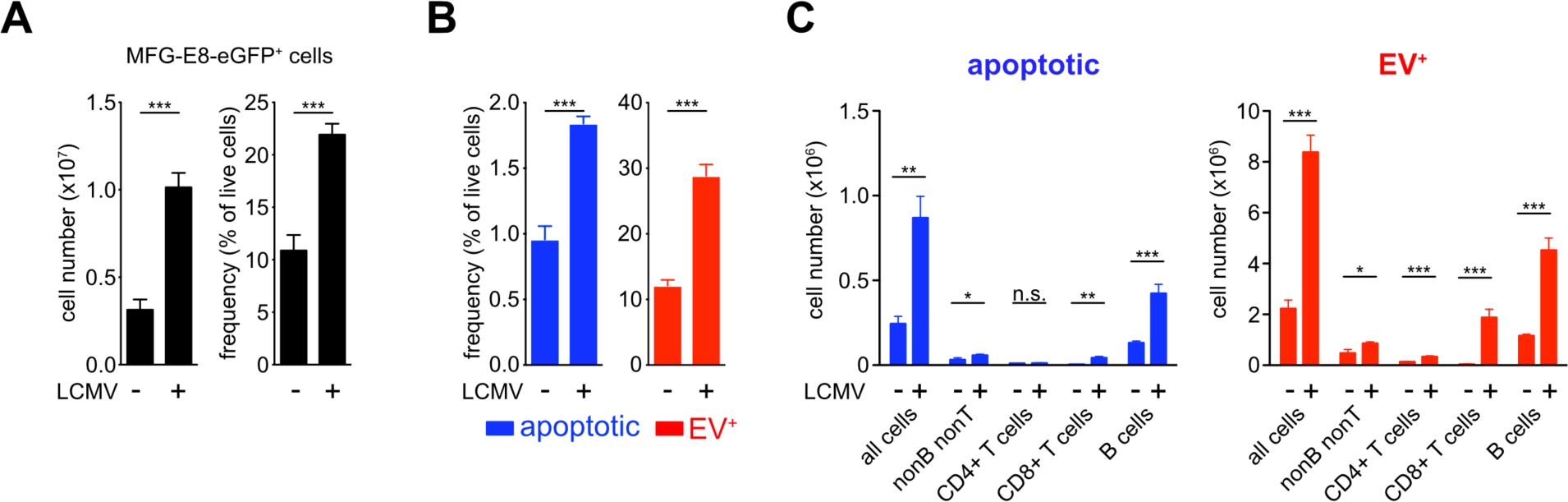
Identification of dying cells and EV^+^ cells during LCMV infection. Non-infected and LCMV_Arm_ (2×10^5^ PFU, i.p.) infected mice were injected with 100µg MFG-E8-eGFP on day 5 post infection. 1h later mice were sacrificed and splenic B, T and non-B/T cell subsets were analyzed by imaging flow cytometry (gating strategy shown in suppl. Fig 4). (A) Bar graphs show total numbers (left) and frequencies (right) of all MFG-E8-eGFP^+^ splenocytes in non-infected and infected mice. (B) Frequencies of MFG-E8-eGFP^+^ apoptotic (blue) and EV^+^ (red) cells were calculated using the CAE. (C) MFG-E8-eGFP^+^ CD19^-^TCRb^-^ nonB/T cells, CD4^+^ and CD8^+^ T cells and CD19^+^ B cells were classified as apoptotic (blue) or EV^+^ (red) using the CAE and their total numbers in the spleen were calculated. Numbers next to the gate show the mean percentage of 3 mice. Bar graphs show averages of 3 mice ± SD. For statistical analysis unpaired Student’s T test was used, asterisks indicate statistical significance. Representative results of 3 independent experiments are shown.

### Detailed analyses of spleen cell populations for apoptotic and EV^+^ cells

Among CD19^+^ B cells, mainly marginal zone (MZ, CD19^+^CD21^hi^CD23^lo^) and follicular (CD19^+^CD21^+^CD23^hi^) B cells showed increased apoptosis, while only follicular B cells showed a significant increase in EV^+^ cells during LCMV-infection (Fig. 6A). MZ B cells showed a very high degree of EV-decoration in both infected and non-infected animals (Fig. 6A). Among the CD19^-^TCRβ^-^ non-B/T cell populations CD11c^+^MHCII^+^ DCs showed slight but significant increases of apoptotic and EV^+^ cells (Fig. 6B).

**Figure 6.**
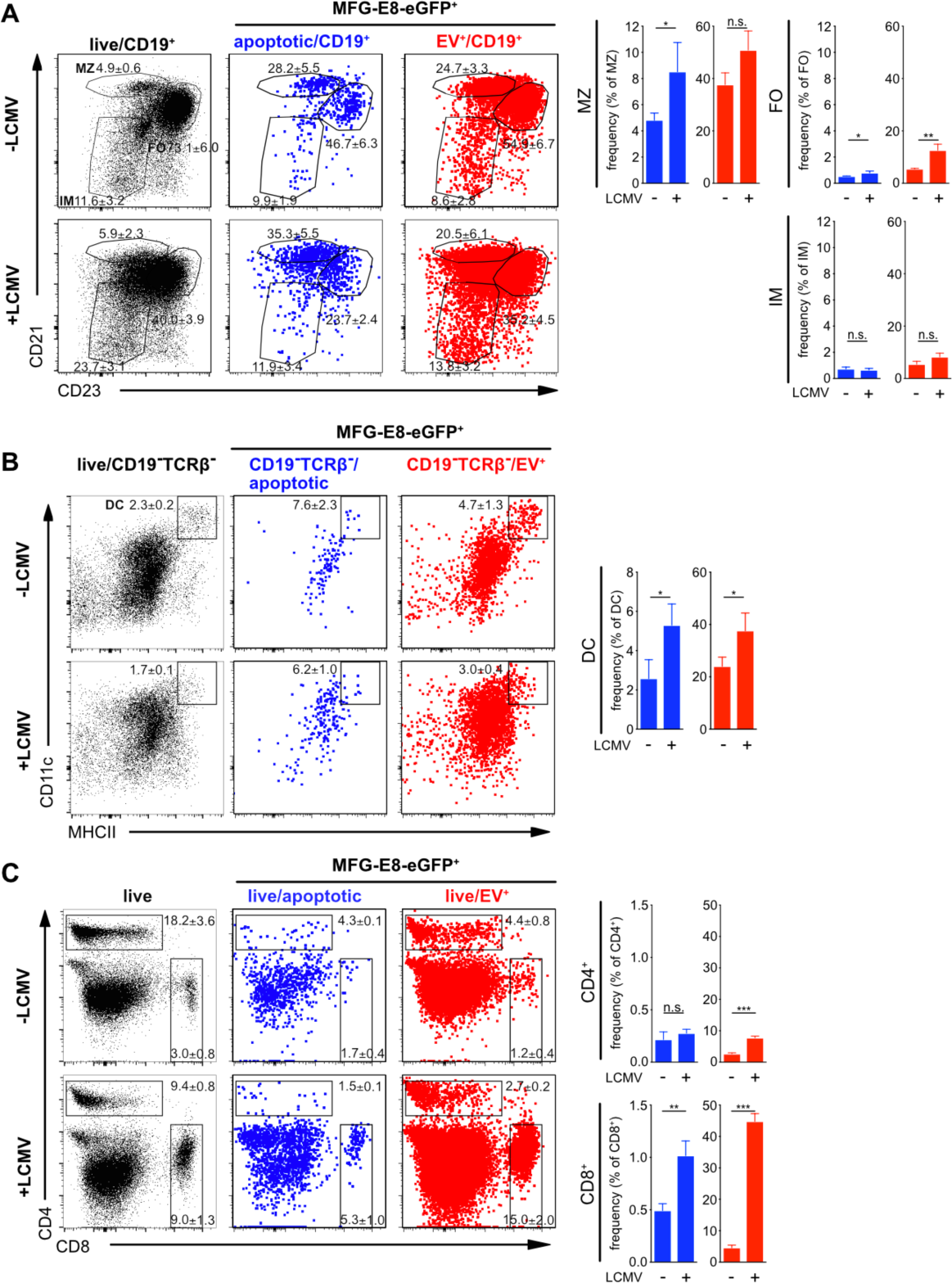
CAE analysis of spleen cell populations during LCMV infection. Non-infected and LCMV_Arm_ (2×10^5^ PFU, i.p.) infected mice were injected with 100µg MFG-E8-eGFP on day 5 post infection. 1h later mice were sacrificed and splenic cell subsets were analyzed by imaging flow cytometry (gating strategy shown in Suppl. Fig 4). (A) MFG-E-eGFP^+^ CD19^+^ B cell subsets (MZ = marginal zone, FO = follicular, IM = immature B cells), (B) CD19^-^TCRβ^-^CD11c^+^MHCII^+^ DCs and (C) CD4^+^ and CD8^+^ T cells were analyzed using the CAE and the percentages of apoptotic and EV^+^ cells determined and depicted in bar graphs. Numbers next to the gate show the mean percentage of 3 mice. Bar graphs show averages of 3 mice ± SD. For statistical analysis unpaired Student’s T test was used, asterisks indicate statistical significance. Representative results of 3 independent experiments are shown.

CD4^+^ and CD8^+^ T lymphocytes play central roles in controlling viral infections (Swain et al., 2012; Wong and Pamer, 2003). Analysis of spleens from LCMV infected mice showed that the frequencies of apoptotic cells increased during viral infection only among CD8^+^ T cells, but not CD4^+^ T cells (Fig. 6C). In contrast, while EV-decoration increased only slightly but statistically significantly also in CD4^+^ T cells, it augmented drastically in CD8^+^ T cells during LCMV infection, when 40-50% of EV^+^CD8^+^ T cells were detected (Fig. 6C). While MZ B cells showed high EV-decoration regardless of LCMV infection, especially CD8^+^ T cells became strongly EV^+^ selectively upon acute LCMV infection. This data shows that Mfge8-eGFP *in vivo* in combination with CAE allows to precisely discriminate apoptotic cells from the vast majority of EV-decorated cells.

### EVs carry markers of exosomes and antigen-presenting cells

We next set out to further characterize cell-attached EVs generated during LCMV-infection *in vivo* using the flow microscopy CAE-pipeline. EVs originating from professional APC like DCs carry exosomal markers such as tetraspanins CD9 and CD63 related to endosomal vesicle trafficking (Buschow et al., 2009; Zitvogel et al., 1998). In addition, they are enriched for DC-markers such as CD86, MHCII and CD54 (Segura et al., 2005). We therefore evaluated next if EV^+^CD8^+^ T cells are positive for these DC-exosome markers. Analysis of naïve CD44^-^CD8^+^ T cells and effector CD44^+^CD8^+^ T cells from the same LCMV-infected animal showed that EVs bound specifically to activated CD44^+^CD8^+^ T cells, but not to resting CD44^-^CD8^+^ T cells (Fig. 7A). We therefore focused on CD44^+^CD8^+^ T cells for further analyses of EVs. We found approximately 80% CD9/CD63^+^, 99% CD54^+^, 78% MHCII^+^ and 98% CD86^+^ CD44^+^CD8^+^ T cells (Fig. 7A). Several of these frequencies and median fluorescence intensities were significantly higher as compared to EV^-^CD44^+^CD8^+^ T cells in infected mice (Fig. 7A), suggesting that decoration with MFG-E8^+^ EVs contributes to the accumulation of APC-markers on effector CD8^+^ T cells.

**Figure 7.**
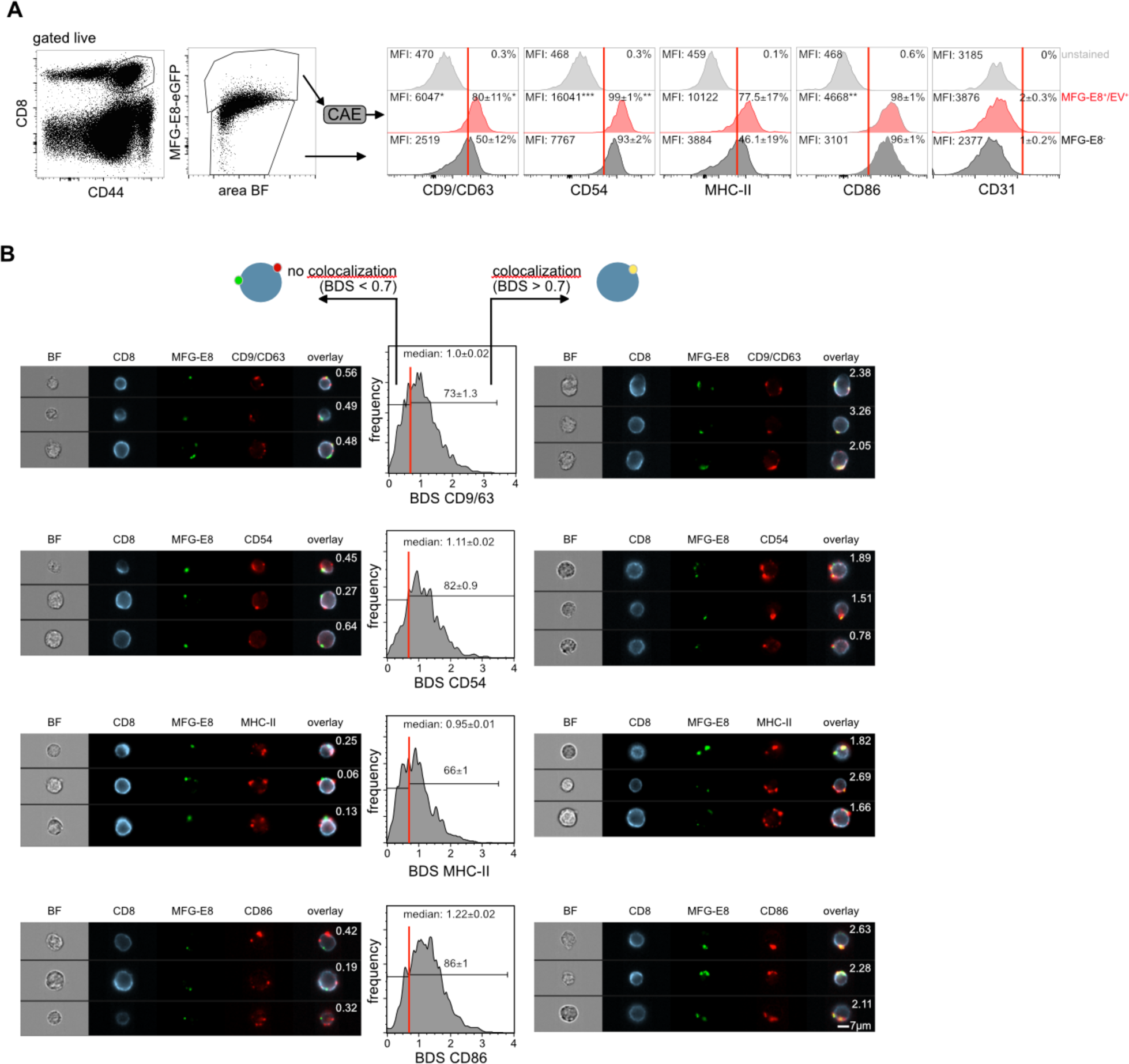
EVs bound to CD8^+^ T cells originate from APCs. (A) Splenic CD8^+^ T cells of LCMV-infected mice (day 5 post infection, n=3) were analyzed for expression the activation marker CD44 and staining for MFG-E8. MFG-E8^-^ and MFG-E8^+^/EV^+^ CD44^+^CD8^+^ T cells were then further analyzed for expression of potential EV-markers CD9/CD63 (combined in one staining), MHCII, CD54, CD86 and CD31. Median fluorescence intensities (MFI) and frequencies of cells positive for these proteins are indicated in the histograms. Asterisks indicate statistical significance. For statistical analysis Student’s T test was used. (B) EV^+^EV-marker^+^ cells were analyzed for their staining pattern of the respective EV-marker. Staining pattern of CD9/CD63, CD54, MHCII and CD86 was similar to MFG-E8^+^ EVs. To determine if EV-marker staining co-localized with MFG-E8-eGFP^+^ EVs overlap of both stainings was quantified using the bright detail similarity (BDS) feature of the IDEAS software. Cells with a BDS < 0.7 did not show any significant co-localization as determined by visual inspection. Cells with a BDS > 0.7 showed at least partial co-localization of MFG-E8 and the respective EV-marker. BDS scores are shown in the representative example images. Median BDS scores and frequency of cells showing co-localization are indicated within histograms.

To determine if these APC-molecules co-localized with MFG-E8-eGFP^+^ EVs or showed a different distribution on the T cell surface, we used the “bright detail similarity” (BDS) feature, a standard co-localization method provided by the IDEAS software. Here, cells with a score <0.7 (Fig. 7B; left panel, BDS <0.7) showed no or weak co-localization, while cells with a score >0.7 (Fig. 7B; right panel, BDS >0.7) showed substantial co-localization. Analysis of the BDS score revealed that on 73% of all EV^+^CD9/63^+^CD8^+^ T cells MFG-E8-eGFP and CD9/63 co-localized on the same EVs (Fig. 7B). We found similarly high frequencies of CD8^+^ T cells with CD54 (82%), MHCII (66%) and CD86 (86%) co-localizing with MFG-E8-eGFP on the same EVs (Fig. 7B). The fact that we also found EVs where MFG-E8 did not colocalize with any of the above markers argues either for the existence of EVs where MFG-E8 staining was absent or below the detection limit, the generation of EVs during organ preparation in absence of sufficient amounts of *in vivo* applied MFG-E8-eGFP or other processes of protein transfer to activated T cells such as trogocytosis (Huang et al., 1999).

It has been shown that many CD31^+^ endothelial cells undergo apoptosis during LCMV infection (Frebel et al., 2012). We therefore also assessed the presence of CD31 on EV^+^ T cells to determine if apoptotic endothelial cells were the origin of T cell-associated EVs. However, we did not find a significant staining for CD31 on T cells (Fig. 7A), nor any CD31^+^ vesicles on T cells (not shown). These results indicate that the great majority of EVs associated with CD8^+^ T cells during LCMV infection are APC-derived exosomes.

## Discussion

Here we report the *in vivo* application of recombinant Mfge8-eGFP for the detection of PS^+^ cells using conventional and image flow cytometry. We show that Mfge8-eGFP binds similar fractions of PS^+^ apoptotic cells *in vitro* as compared to the more widely used Annexin V. However, PS-binding of Mge8-eGFP is calcium independent and works after *in vivo* injection. Surprisingly, we found the majority of PS^+^ cells in irradiated and LCMV-infected mice being not apoptotic, but alive and decorated with PS^+^ EVs. Upon development of a deep learning autoencoder, we can faithfully separate both, true apoptotic cells and EV-decorated cells for further analyses.

The detection of cell death in tissues *in vivo* is challenging. Therefore, dead cell analyses are mostly performed *ex vivo* with single cell suspensions derived from the organs of interest after biopsies or sacrifice of experimental animals. However, this procedure exposes cells to additional stress factors such as shear forces by tissue homogenization, enzymes, temperature and pH changes, salt compositions of working solutions and many more. Hence, tissue preparation for dead cell analysis *ex vivo* might artificially increase cell death rates due to handling. In addition, most methods to measure apoptotic cells have intrinsic restrictions, adding to their imprecision of analyzing cell death. For example, labeling of fragmented DNA by TUNEL (terminal deoxynucleotidyl transferase dUTP nick end labeling) mostly detects late stage apoptotic cells only (Negoescu et al., 1996), which are very rapidly cleared *in situ*, as most DNA fragmentation of dying cells occurs inside phagocytes (Odaka and Mizuochi, 2002). Measuring the active form of caspase-3 using fluorescent substrates is not completely specific, as caspase-3 is also activated independently of cell death in certain cell types (McComb et al., 2010). The *ex vivo* staining of cells for surface phosphatidylserine (PS) using Annexin V has the disadvantage to require high Ca^2+^-levels, precluding it from most *in vivo* applications and interfering with many other downstream applications (van Engeland et al., 1998). MFG-E8, also known as lactadherin, also binds to PS on apoptotic cells and enhances their engulfment by phagocytes (Hanayama et al., 2002). Translocation of PS to the outer membrane not only occurs during apoptosis, but also during the formation of microvesicles (Hugel et al., 2005; Martinez and Freyssinet, 2001) and exosomes (Thery et al., 2002), allowing detection of PS^+^ EVs by MFG-E8-eGFP.

Although we could detect many MFG-E8^+^ cells, true apoptotic cells were rare in spleens and BM of control mice, while the great majority were EV-decorated live cells. Due to their great morphological variability, a reliable discrimination was not possible using fluorescent intensity of the MFG-E8 signal only. Also, a combination of two features that worked well on manually selected apoptotic and EV-decorated cells failed in more complex samples. Only *in vivo* administered MFG-E8-eGFP in combination with imaging flow cytometry and a deep learning approach using a CAE-RF allowed us to reliably classify the MFG-E8^+^ cells into PS^+^ apoptotic or PS^+^ EV-decorated cells. Previous reports showed substantial death among T cells during the early phase (day 2-4) of infection LCMV infection (Bahl et al., 2010), which is dependent on type I interferon production (Crouse et al., 2015) and even affects nonspecific bystander cells (Jiang et al., 2003). In contrast, we could identify only low numbers of dying cells in acutely LCMV infected mice. The reason for this discrepancy could lie in the different methods to identify dying cells or could be due to different time points of analysis.

Another striking finding is the dramatic increase in cells that bind EVs. On the one hand these vesicles could be virus-containing particles infecting new cells, or apoptotic bodies reflecting the increased amount of cell death. However, given the wide range of immunoregulatory functions of EVs (Robbins and Morelli, 2014), the increase in EVs could also be a consequence of the ongoing immune response. The fact that especially B cells and activated CD8^+^ T cells bind these EVs, supports this idea. It has been shown previously that activated DCs secrete MHC class II containing exosomes, which bind to activated CD4^+^ T cell via LFA-1 (Nolte-’t Hoen et al., 2009) and could play a role in T cell activation (Buschow et al., 2009). Evidence for their strong immunostimulatory function came from early exosome studies demonstrating the capability of tumor-antigen bearing exosomes secreted from DCs to trigger T-cell dependent anti-tumor responses (Zitvogel et al., 1998). However, in our study CD4^+^ T cells were not as strongly EV-decorated as compared to CD8^+^ T cells in LCMV-infected mice. Previous reports showed that vesicles derived from DCs were able to stimulate CD8 T cells *in vitro* (Kovar et al., 2006) and transfer exogenous antigen to DCs for CD8^+^ (Winau et al., 2006) and CD4^+^ T cell priming (Montecalvo et al., 2008).

Approximately half of all CD8^+^ T cells were EV-decorated during LCMV infection. This could indicate that either those T cells were targeted by EVs produced by other cells, or that we detected nascent EVs produced by T cells themselves. Both scenarios are possible. It has been shown that TNFα-containing exosomes were able to delay activation-induced cell death in T cells (Zhang et al., 2006). On the other hand, T cells release EVs constitutively and EV secretion is enhanced by TCR triggering (Blanchard et al., 2002; van der Vlist et al., 2012), which causes increased intracellular calcium levels for enhanced EV-production (Savina et al., 2003). Moreover, EVs from CD8^+^ T cells may also contain granzyme and perforin (Peters et al., 1991) and can inhibit antigen presentation and survival of DCs to downmodulate immune responses in mouse models of cancer and diabetes (Xie et al., 2010). In addition, microvesicles budding from the immunological synapses of CD4 T cells *in vitro* do contain TCR, which may transfer signals to B cells expressing cognate peptide MHCII (Choudhuri et al., 2014). However, our finding that MFG-E8+ EVs carry DC-markers such as MHCII, CD86, CD54 and tetraspanins rather argue for the exosomal APC-origin of these EVs.

Also, many B cells carried vesicles, even in non-infected mice. EVs can be a source of native, unprocessed antigen (Qazi et al., 2009) and in the case of virus infections they could carry intact viral proteins (Nolte-’t Hoen et al., 2016) for recognition by cognate B-cell receptors causing B cell activation. In addition, previous studies described B cells becoming Annexin V positive, without undergoing apoptosis (Dillon et al., 2001). While the authors described that PS-exposure was selective for B cells upon their IgM-mediated positive selection, they did not determine, if PS-exposure was cell-intrinsic or by EV-decoration.

Further studies are necessary to determine if EV binding is restricted to certain zones of lymphatic organs, such as the MZ, where blood is filtered. Cells in the MZ, such as MZ B cells, CD11c^+^CD11b^-^ DCs and activated T cells locate to this region and could therefore be preferentially exposed to EVs. However, the fact that also follicular B cells are EV-decorated argues against this possibility.

To our knowledge, this is the first report that identifies cell subsets binding naturally occurring EVs *in vivo. In vivo* staining of dying cells and EV-decorated cells using Mfge8-eGFP is a valuable tool, not only to reliably identify cells that undergo cell death in different pathological conditions, but also to clarify the function of EVs on different cell types in various tissues under normal and pathological conditions, such as viral infections, autoimmunity and cancer.

## ACKNOWLEDGEMENTS

We acknowledge the Core Facility Flow Cytometry at the Biomedical Center, Ludwig-Maxmilians-Universität München, for providing the ImageStream^X^ MKII imaging flow cytometer. N.K.C., A.L., T.K. were supported by a Deutsche Forschungsgemeinschaft (DFG, German Research Foundation) fellowship through the Graduate School of Quantitative Biosciences Munich (QBM) and N.K.C. is supported additionally through the School of Life Sciences Weihenstephan, Technical University of Munich, Germany. F.J.T. acknowledges financial support by the Graduate School QBM, the DFG within the Collaborative Research Centre (CRC) 1243 (Subproject A17), by the Helmholtz Association (Incubator grant sparse2big, grant # ZT-I-0007), by the BMBF (grant# 01IS18036A and grant# 01IS18053A) and by the Chan Zuckerberg Initiative DAF (advised fund of Silicon Valley Community Foundation, 182835). T.B. is supported by the DFG CRC 1054 (TP B03) and Graduate School QBM. This work was funded by the DFG under Germany’s Excellence Strategy within the framework of the Munich Cluster for Systems Neurology (EXC 2145 SyNergy – ID 390857198) to M.K.

## AUTHOR CONTRIBUTIONS

N.K.C. and F.J.T. developed the deep learning model and the data analysis pipeline. T.B. and J.K. planned experiments and wrote the paper. L.R., A.L., A.F.-A.K. and T.K. performed experiments. M.SCH. and M.S. provided electron microscopy expertise.

