## Supplemental files for "*In vivo* identification of apoptotic and extracellular vesicle-bound live cells using image-based deep learning"

### Supplemental Information

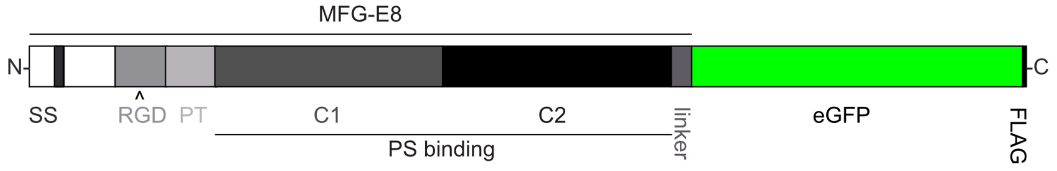


Supplemental Figure 1

**Supplemental Figure 1: MFG-E8-eGFP fusion protein**

Enhanced green fluorescent protein (eGFP) was fused to the C-terminus of full-length murine MFG-E8, separated by a 15aa helical linker (linker). To aid purification, a FLAG-tag was added to the C-terminus of eGFP. PT, prolin-threonine rich domain; RGD, RDG-motif; SS, signal sequence; N, N-terminus; C, C-terminus; C1, C1-domain; C2, C2-domain; PS, phosphatidylserine;

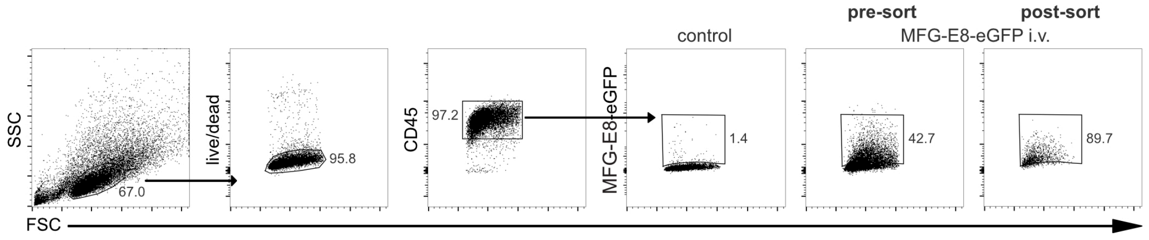


Supplemental Figure 2

**Supplemental Figure 2: Sorting strategy for MFG-E8$^+$ splenocytes**

$LCMV_{Arm}$ infected mice (day 5 post infection) were injected with 100µg MFG-E8-eGFP i.v. 1h prior to sacrifice. Live/dead$^-$CD45$^+$MFG-E8-eGFP$^+$ splenocytes were sorted. Sort purity of MFG-E8-eGFP$^+$ cells was approx. 90%. Cells were then analyzed for the presence of extracellular vesicles by TEM.

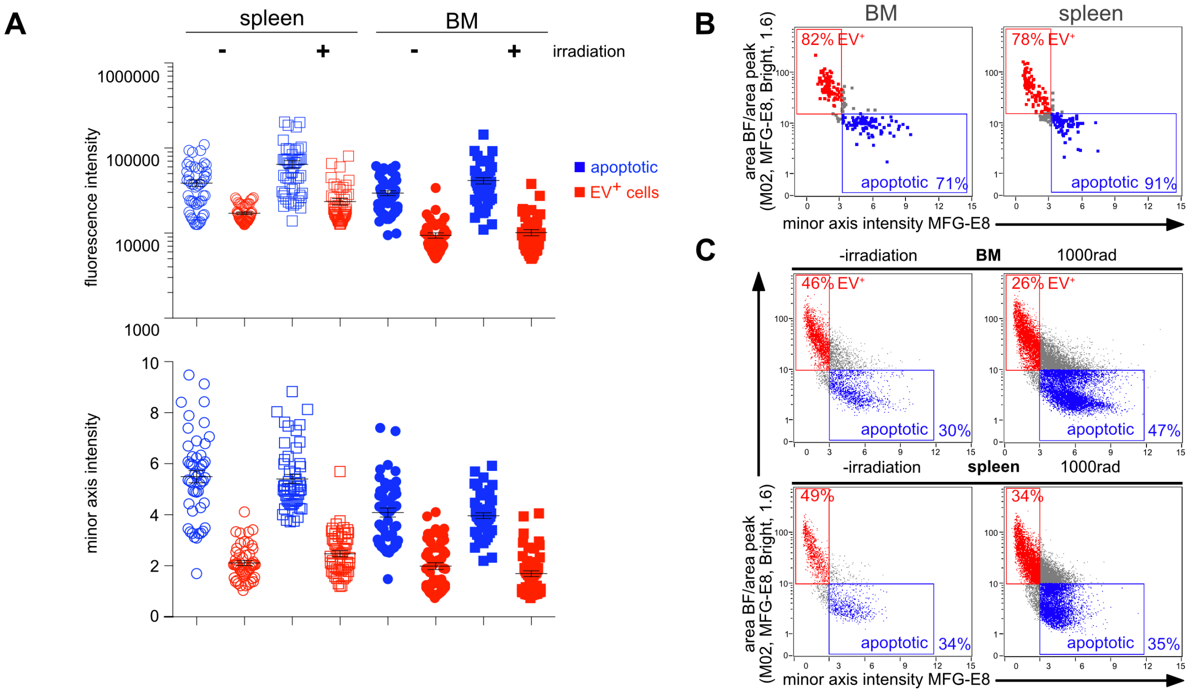


**Supplemental Figure 3: Comparison of MFG-E8-eGFP MFI and minor axis intensity and IDEAS gating strategy.** (A) Mean MFG-E8-eGFP fluorescence intensities (top) and MFG-E8-eGFP minor axis intensities (bottom) of 200 manually selected apoptotic cells and 200 EV$^+$ cells from irradiated and non-irradiated BM cells and splenocytes are shown. Using minor axis intensities resulted in a good separation of apoptotic and EV$^+$ cells in both spleen and BM, while using the MFI of MFG-E8-eGFP did not result in a good separation of apoptotic and EV$^+$ cells in BM. (B) 200 apoptotic (blue) and 200 EV$^+$ cells (red) were manually selected from non-irradiated and irradiated spleen and BM samples. Dot plots show their MFG-E8-eGFP minor axis intensity values and the ratio of the BF area and the area of the MFG-E8-eGFP signal. Apoptotic cells were defined as having a MFG-E8-eGFP minor axis intensity value >3 and an area ratio <10, EV$^+$ cells as having a MFG-E8-eGFP minor axis intensity <3 and an area ratio >10. Percentages show the frequency of manually selected cells that were correctly classified by the IDEAS features. (C) Apoptotic cells (blue) and EV$^+$ cells (red) from total spleen and BM from irradiated and non-irradiated mice were quantified using the BF:MFG-E8-eGFP area ratio and the MFG-E8-eGFP minor axis intensity. Uncategorized cells are shown in gray.

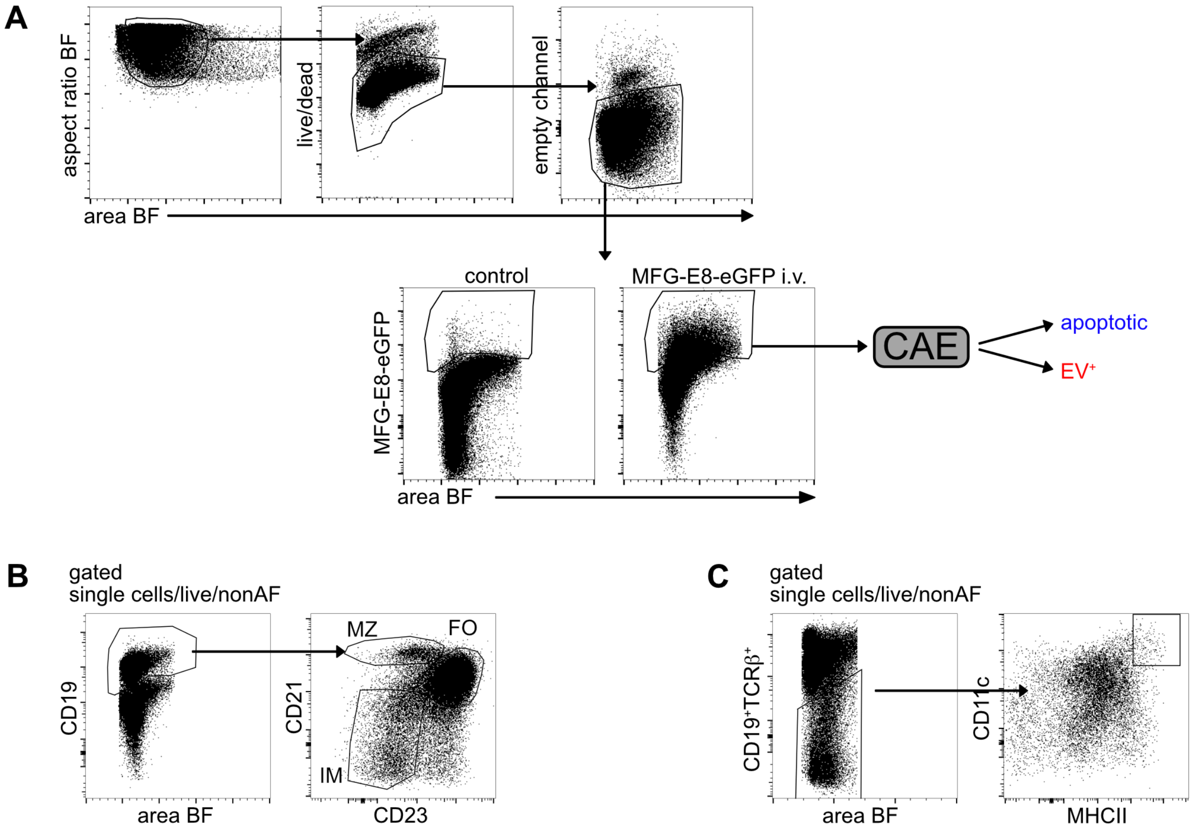


**Supplemental Figure 4: Flow cytometry gating strategies.**

(A) General gating strategy for MFG-E8$^+$ apoptotic and EV$^+$ cells is shown. First, single cells are gated using the aspect ratio and the area of the BF signal. Then necrotic cells are removed by gating live/dead viability dye negative cells (live). If required, autofluorescent cells were removed by using an empty channel (channel 4). Then MFG-E8-eGFP$^+$ cells were gated using a negative control (PBS injection). MFG-E8-eGFP$^+$ cells were sorted into apoptotic and EV$^+$ cells using a CAE. (B) Gating strategy for different B cell subsets: First, all B cells were gated using CD19. Then different subsets were gated based on their CD21 and CD23 expression (MZ = marginal zone, CD21$^+$CD23$^-$; FO = follicular, CD21$^+$CD23$^+$; IM = immature B cells, CD21$^-$CD23$^-$). (C) Gating strategy to analyze DCs: First, non-lymphocytes were gated as CD19$^-$/TCR$\beta^-$ cells. Then DCs were gated as CD11c$^+$MHCII$^+$. (D) For analysis of NFAT2 translocation MFG-E8$^-$ and MFG-E8$^+$ cells were gated. Then MFG-E8$^-$CD8$^+$CD44$^+$ and MFG-E8$^+$CD8$^+$CD44$^+$ populations were determined. Of these, Draq5$^+$NFAT2$^+$ cells were selected. To determine correct gating of NFAT2$^+$ cells, a negative control without NFAT2 antibody was used.

| anti-mouse antibodies | | | |
|---|---|---|---|
| **reactivity/ staining reagent** | **conjugate** | **clone** | **vendor** |
| CD11b | APC | M1/0 | eBioscience |
| CD11c | PE/Cy7 | N418 | BioLegend |
| CD19 | APC/eFluor780 | eBio1D3 | eBioscience |
| CD19 | PE/Cy7 | 6D5 | BioLegend |
| CD19 | APC | 1D3 | BD Biosciences Pharmingen |
| CD21/35 | APC/Cy7 | 7E9 | BioLegend |
| CD23 | PE/Cy7 | B3B4 | BioLegend |
| CD31 | AF647 | MEC13.3 | BioLegend |
| CD4 | APC | RM4-5 | BioLegend |
| CD44 | APC/Cy7 | IM7 | BioLegend |
| CD44 | Pacific Blue | IM7 | BioLegend |
| CD45 | AF647 | 30-F11 | BioLegend |
| CD51 | PE | RMV-7 | Biolegend |
| CD54 | AF647 | YN1/1.74 | Biolegend |
| CD61 | PE | 2C9.G2 | Biolegend |
| CD62L | PE | MEL-14 | eBioscience |
| CD63 | APC | NVG-2 | eBioscience |
| CD8a | PE/Cy7 | 53-6.7 | BioLegend |
| CD86 | PE | GL1 | eBioscience |
| CD9 | APC | eBioKMC8 | eBioscience |
| cleaved Cas8 | purified | D5B2 | Cell Signaling |
| GFP | FITC | polyclonal | Abcam |
| MHC-II (I-A/I-E) | PE/Cy7 | M5/114.15.2 | eBioscience |

| other staining reagent | | |
|---|---|---|
| **reactivity/ staining reagent** | **conjugate** | **vendor** |
| AnnexinV | Cy5 | Abcam |
| live/dead violet | | Thermo Fisher |
| PKH26 | | Sigma Aldrich |

**Suppl. Table 1: Staining reagents and antibodies used in this study.**
